# Tuning the SMC: efficient simulation and the structure of ARGs

**DOI:** 10.64898/2026.09.01.748538

**Authors:** Hossameldin Loay, Gertjan Bisschop, Derek Setter, Abdelmajid Omarjee, Jerome Kelleher, Konrad Lohse

**Author notes:** Joint first author (Hossameldin Loay, Gertjan Bisschop). Joint senior author (Jerome Kelleher, Konrad Lohse).

## Abstract

Sequentially Markovian Coalescent (SMC) models are a central element of contemporary population genetics, underlying many inferential methods. While the SMC has been shown to closely approximate the canonical Coalescent with Recombination (CwR) in terms of lowdimensional, two-locus summaries, its effects on the deeper structural properties of Ancestral Recombination Graphs (ARGs) are less well understood. Here, we define a general SMC approximation, SMC(*k*), in which a single parameter *k* controls the physical scale over which common-ancestor events between non-overlapping ancestral segments are permitted. The model encompasses the standard SMC and SMC^*′*^ as special cases and converges to the CwR as *k* increases, providing a tunable trade-off between computational efficiency and fidelity to the full recombination process. Using recently developed summaries of ARG structure, we show that SMC approximations systematically truncate the persistence of ancestral haplotypes across the genome, despite preserving marginal coalescent properties, and that increasing *k* progressively recovers this long-range ancestral structure. We implement the SMC(*k*) in `msprime` and show that, for small samples, it makes whole-chromosome simulation in species with large population-scaled recombination rates several orders of magnitude faster than the CwR. Finally, we use SMC simulations for chromosome-scale parametric bootstrapping of demographic inference and find that the SMC^*′*^ captures uncertainty in SFS-based estimates remarkably well, with only modest changes as *k* increases despite substantial differences in long-range ARG structure. Thus, the importance of SMC approximation error depends strongly on which properties of ancestry are relevant to the downstream analysis.

## Introduction

The coalescent with recombination (CwR) is the canonical mathematical framework for modelling the ancestry of recombining sequences backwards in time (Hudson 1983; Griffiths and Marjoram 1997). Although the CwR captures the full stochastic process of recombination and common ancestry, its complexity has long made both inference and simulation challenging. McVean and Cardin (2005) introduced the Sequentially Markovian Coalescent (SMC) as an approximation in which the sequence of local genealogies forms a Markov process along the genome. Viewed backwards in time, this approximation can be understood as restricting which ancestral lineages are permitted to coalesce: under the standard SMC, common-ancestor events occur only between lineages carrying overlapping ancestral material. The SMC^*′*^ refinement additionally permits adjacent ancestral segments to merge (Marjoram and Wall 2006). The SMC and SMC^*′*^ have since become standard models in population genetics and underpin a wide range of inferential methods (Peede et al. 2026). SMC approximations preserve the marginal distribution of local genealogies, but necessarily alter their dependence along the genome. How important these differences are has classically been assessed using low-dimensional summaries of statistical association. Early work compared linkage disequilibrium in simulations (e.g. McVean and Cardin 2005), while later analytical results showed that the SMC^*′*^ provides an accurate approximation to the joint distribution of pairwise coalescence times at two loci (Wilton *et al*. 2015). However, pairwise and two-locus summaries capture only part of the dependence encoded by a complete Ancestral Recombination Graph (ARG), and it is less clear how SMC approximations affect the persistence of ancestral haplotypes and other long-range structural properties of ARGs.

Recent interest in ARG inference (Brandt *et al*. 2024; Lewanski *et al*. 2024; Nielsen *et al*. 2024; Wong *et al*. 2024) has motivated new ways of describing and comparing the structure of inferred and simulated ARGs. In particular, summaries based on the genomic spans associated with ancestral nodes and on haplotype-based ARG similarity provide information about ancestry that is not captured by local trees or pairwise linkage statistics (Ignatieva *et al*. 2025; Fritze *et al*. 2026). These developments make it possible to revisit the accuracy of the SMC family at the level of the ARG itself. This is increasingly relevant as population-genetic analyses make greater use of long-range haplotype and genealogical structure (e.g. Pope *et al*. 2026), for which agreement in marginal or two-locus distributions need not imply agreement in the features of ancestry relevant to downstream inference.

An early application of SMC approximations was to provide faster simulations. Early CwR simulators such as `ms` (Hudson 2002) could not efficiently generate whole chromosomes under human-like parameters, motivating approximate simulators based on the SMC and related models (Marjoram and Wall 2006; Chen et al. 2009; Staab et al. 2015). The introduction of `msprime` (Kelleher et al. 2016) largely removed this computational advantage for human-like parameter regimes through an efficient implementation of Hudson’s CwR simulation algorithm. Over the past decade, `msprime` has developed into a general platform for ancestry simulation (Ragsdale et al. 2020; Baumdicker *et al*. 2022; Kelleher and Lohse 2020), with support for models including the discrete-time Wright–Fisher process (Nelson et al. 2020), explicit pedigrees (Anderson-Trocmé et al. 2023), and hybrid forward- and backwards-time simulations (Kelleher et al. 2018; Haller et al. 2018; Gopalan et al. 2025). Its integration with `tskit` (Jeffery et al. 2026) also provides efficient access to the resulting ARGs. Nevertheless, the computational cost of exact CwR simulation grows quadratically with population-scaled recombination rate *ρ* (Baumdicker et al. 2022), and so whole-chromosome simulation can be prohibitively expensive for species with large *ρ*, such as *Drosophila melanogaster*.

Here, we define a generalised SMC approximation, SMC(*k*), in which the parameter *k* controls the maximum physical separation over which common-ancestor events between nonoverlapping ancestral segments are permitted. The standard SMC and SMC^*′*^ models arise as special cases, while increasing *k* progressively approaches the full CwR. Related approaches have previously been used to trade approximation against simulation speed (Chen et al. 2009; Shlyakhter et al. 2014; Staab et al. 2015); here, we use this generalisation both as an efficient simulation model and as a controlled framework for characterising what aspects of ARG structure are lost under SMC approximations. We implement the SMC(*k*) efficiently in `msprime` v1.4 and show that it enables whole-chromosome simulation in high-recombination parameter regimes that are prohibitively expensive under the CwR. Using recently developed summaries of ARG structure, we show that low-*k* approximations truncate the persistence of ancestral haplotypes along the genome, with CwR-like longrange structure progressively recovered as *k* increases. Finally, we illustrate how these computational gains make previously impractical analyses possible by using chromosome-scale SMC simulations to quantify uncertainty in demographic inference through parametric bootstrapping in *D. melanogaster*. Despite substantial differences in long-range ARG structure, we find that the SMC^*′*^ captures uncertainty in SFS-based demographic estimates remarkably well, illustrating that the practical importance of SMC approximation error depends on the properties of ancestry relevant to the downstream analysis.

## Results

### The SMC(k) model

The SMC models the ancestry of a sample by assuming a Markovian process along the genome, such that the local genealogy at any position depends only on the genealogy immediately to its left (McVean and Cardin 2005). Backwards in time, the standard SMC has an equivalent formulation in which common ancestor events are restricted to pairs of lineages carrying overlapping ancestral material (McVean and Cardin 2005; Bisschop et al. 2026). This restriction excludes common ancestor events that generate trapped, non-ancestral material (Wiuf and Hein 1999), causing the SMC to overestimate the number of independent ancestors contributing to a sample and to underestimate long-range linkage disequilibrium (Marjoram and Wall 2006). For example, under the CwR, two lineages created by a recombination event may subsequently coalesce back together before further events alter their ancestry. The SMC^*′*^ restores these “diamonds” (Rasmussen *et al*. 2014) by additionally permitting common ancestor events between exactly adjacent, but non-overlapping, ancestral segments (Marjoram and Wall 2006). This backwards-time formulation suggests a natural family of intermediate approximations in which common ancestor events are permitted over a genomic scale.

We define the SMC(*k*) by associating each ancestral lineage with the genomic “hull” of the ancestral material that it carries, spanning its leftmost to rightmost ancestral segment. A common ancestor event between lineages *a* and *b* is permitted only when the physical distance between their hulls is at most *k* (Fig. 1). Thus, *k* determines the maximum genomic scale over which separated ancestral material can be brought together by a common ancestor event. Importantly, the SMC(*k*) modifies only which common ancestor events are permitted; recombination events occur at the same local rates as under the CwR. On a genome with discrete coordinates and a sequence length of *L* base pairs, SMC(0) is equivalent to the standard SMC, and SMC(1) to the SMC^*′*^, while SMC(*L*) recovers the full CwR. Increasing *k* therefore gives a nested sequence of models in which progressively more of the common ancestor events possible under the CwR are retained. Consequently, increasing *k* permits progressively longer spans of trapped non-ancestral material (Fig. 1). Related approaches to controlling the extent of the SMC approximation have previously been implemented in MaCS (Chen *et al*. 2009), scrm (Staab *et al*. 2015), and Cosi2 (Shlyakhter *et al*. 2014).

**Figure 1.**
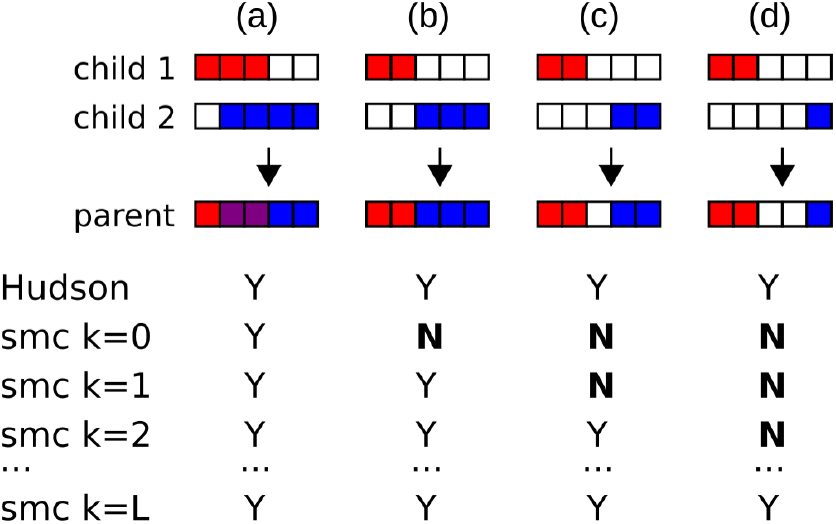
The SMC(k) model restricts common ancestry events. We indicate which of the possible common ancestor events are permitted for various choices of *k*. All scenarios shown in columns (a)–(d) are allowed under the CwR. For *k* = 0 (SMC), common ancestry (purple) is permitted only when the lineages carry overlapping ancestral material (red and blue; column (a)). For *k* = 1 (SMC’), common ancestry may also occur if adjacent intervals are present (columns (a) and (b)). For *k* > 1, common ancestry events that result in trapped non-ancestral material are permitted (columns (c) and (d)). The maximum span of any such interval is *k* − 1, and when *k* = *L*, the full sequence length, the model converges to the CwR (column (d)).

We implemented the SMC(*k*) in `msprime` v1.4 by extending its existing backwards-time CwR simulation algorithm (Kelleher *et al*. 2016), using a similar approach to Cosi2 (Shlyakhter *et al*. 2014). In addition to the linked list of ancestral segments maintained for each lineage, the SMC(*k*) implementation records the hull of each extant lineage. Hull endpoints are maintained in ordered search structures, allowing the simulator to identify eligible common-ancestor pairs without explicitly testing all possible lineage pairs after each event. A separate cumulative index stores, for each lineage, the number of eligible common-ancestor partners and allows common-ancestor events to be sampled efficiently. The sum of these counts determines the rate of common ancestor events, and when such an event occurs a pair is sampled uniformly from the set of eligible pairs. Recombination and common ancestor events can change the extent of the ancestral material carried by a lineage, in which case its hull and the corresponding pair counts are updated. Thus, relative to Hudson’s CwR algorithm, the principal additional cost of the SMC(*k*) is maintaining the search indexes that identify eligible common-ancestor pairs.

This additional bookkeeping leads to a different computational regime from the standard CwR algorithm. For modest sample sizes and very large population-scaled recombination rates, the SMC(*k*) can provide substantial gains over the CwR. For example, simulating two diploid samples for the entirety of *Drosophila melanogaster* chromosome 2R under SMC(1) is approximately three orders of magnitude faster than under the CwR (Fig. 2). This makes chromosome-scale simulations feasible in a parameter regime in which exact CwR simulations are prohibitively expensive. The advantage is not universal, however. Over the range tested, execution time for SMC(1) scales approximately linearly with sample size, whereas the corresponding CwR simulations scale approximately logarithmically (Supplementary Figs. S1, S2). The computational advantage of the SMC(*k*) also diminishes as *k* increases. Consequently, the SMC(*k*) is most useful for long sequences with large populationscaled recombination rates and relatively modest sample sizes; for large samples or sufficiently large *k*, the standard CwR implementation can be more efficient.

**Figure 2.**
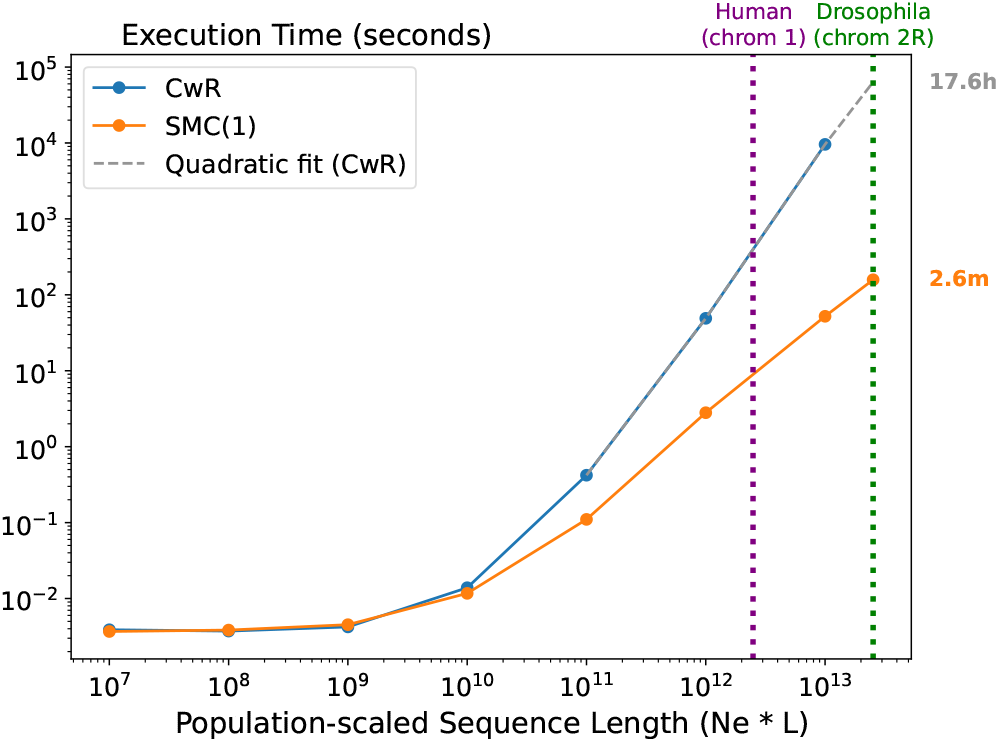
The SMC(*k*) enables whole chromosome simulations in the *Drosophila* parameter regime. Execution time for simulations under the CwR and the SMC(1) model for a sample of *n* = 10 diploid sequences and assuming different population-scaled sequence length for chromosome 1 in humans (10^4^ *×* 249*Mbp*) and chromosome 2R in *D. melanogaster* (10^6^ *×* 25.3*Mbp*).

### Long-range ARG structure under SMC approximations

The SMC^*′*^ is widely regarded as a highly accurate approximation to the CwR, supported by the close agreement between the two models in the joint distribution of pairwise coalescence times at two loci (Wilton *et al*. 2015; Marjoram and Wall 2006). Expectations for standard two-locus summaries of linkage disequilibrium can be expressed in terms of the covariance in pairwise coalescence times (McVean 2002; Lohse *et al*. 2011), but such summaries capture only part of the dependence encoded by a complete ARG. In particular, close agreement in two-locus quantities does not establish whether SMC approximations preserve the long-range persistence of ancestry across the genome. To characterise these differences directly, we therefore consider structural properties of the nodes contained in ARGs generated under the SMC(*k*) and the CwR.

Each node in an ARG corresponds to a unique genetic ancestor of the sample, and the genomic material associated with a node can be divided into three categories (Fig. 3). The binary span of a node is the total genomic span over which an ancestor contributes to two child nodes. Analogously, the unary span is the total genomic span over which an ancestor has a single child node (Wong *et al*. 2024). Unary structure captures the persistence of ancestral lineages across neighbouring local genealogies and is increasingly recognised as an important component of overall ARG structure (Bisschop *et al*. 2026; Fritze *et al*. 2026). Finally, trapped non-ancestral material consists of genomic intervals lying between binary and/or unary segments associated with the same node. A single common ancestor node may therefore contribute to more than one span category. For example, node 5 in Fig. 3 is binary over two genomic segments separated by trapped non-ancestral material, whereas node 7 is binary over the same two segments but unary across the intervening interval. Together, these span measures describe how ancestral lineages persist across the genome in ways that are not captured by the sequence of local trees alone.

**Figure 3.**
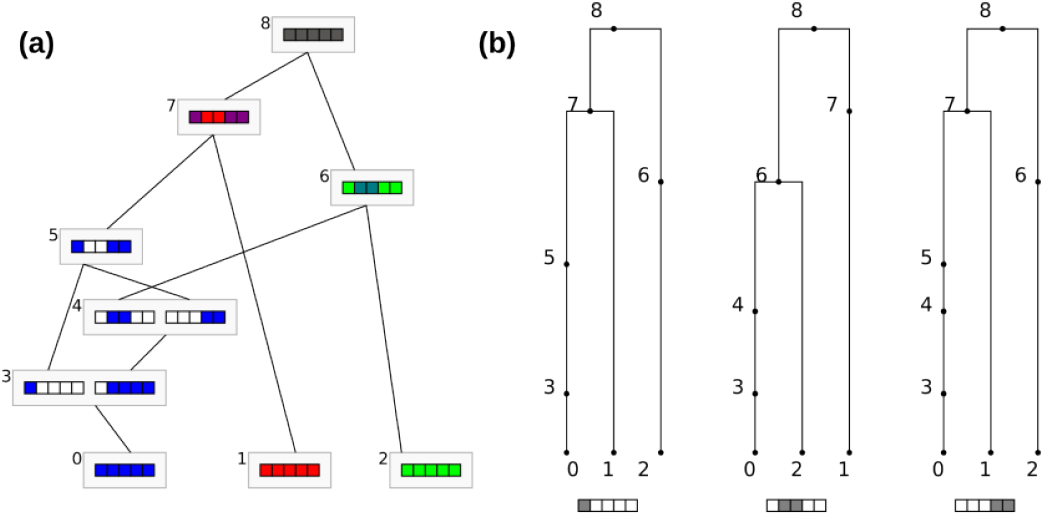
The ancestry of a sample of three haplotypes as a) a graph and b) sequence of local trees. In the ARG, we see that nodes 3 and 4 both have two parents, meaning they are recombinant nodes. These recombination events result in three distinct correlated trees along the genome. The common ancestor node 5 has trapped non-ancestral material. It is (locally) unary in the left and right trees but absent in the middle tree. Subsequent coalescence in node 7 results in a node that is binary for the left and right trees, but unary in the middle tree. This type of event is possible only under the CwR or the SMC(k) with k ≥ 3.

We simulated ARGs under the SMC(*k*) for a range of *k* values and compared the resulting node spans with those generated under the CwR. Across the scaled recombination rates considered, low-*k* approximations produce systematically shorter unary spans than the CwR (Fig. 4). Binary spans show the same qualitative pattern (Supplementary Fig. S3), and similar results are obtained for larger samples. These differences indicate that ancestral lineages persist across shorter genomic distances under the SMC(*k*). As *k* increases, both unary and binary spans progressively approach those observed under the CwR, while increasingly long intervals of trapped non-ancestral material become possible. Thus, common ancestor events between separated genomic segments, although contributing relatively little to pairwise coalescence-time correlations, make an appreciable contribution to the long-range persistence of ancestral material in a complete ARG.

**Figure 4.**
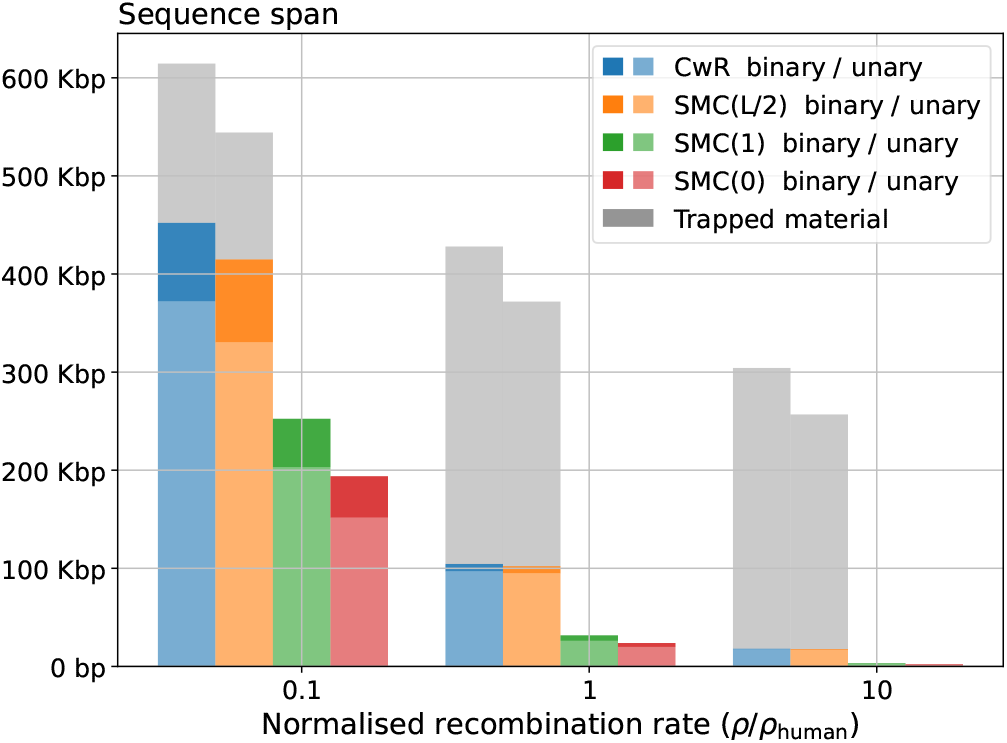
The distribution of ancestral material differs between the SMC’ and CwR models. Stacked bars show the average sequence span per node for unary nodes (lighter colours), binary nodes (darker colours), and trapped non-ancestral material (grey) across human-scaled recombination rates for different models. ARGs that arise from the SMC’ and standard SMC model do not contain trapped non-ancestral material by definition. The simulated chromosome is of 1 Mbp length.

We next compared the overall ancestral structure of ARGs using “matched span”, a similarity measure implemented in *tscompare* (Fritze et al. 2026). We normalised matched-span similarity by the baseline similarity expected between independently simulated CwR ARGs. The fraction of CwR ancestral structure matched by SMC(*k*) ARGs increases with *k*, approaching the CwR baseline as progressively more common ancestor events are permitted (Fig. 5). In contrast, the reciprocal comparison shows little dependence on *k*: ancestral structure present in SMC(*k*) ARGs is generally also represented in CwR ARGs (Supplementary Fig. S5). Together, these results are consistent with the SMC(*k*) generating a restricted subset of the ancestral structures produced under the CwR, with low-*k* approximations primarily removing long-range ancestral relationships rather than introducing qualitatively different structures.

**Figure 5.**
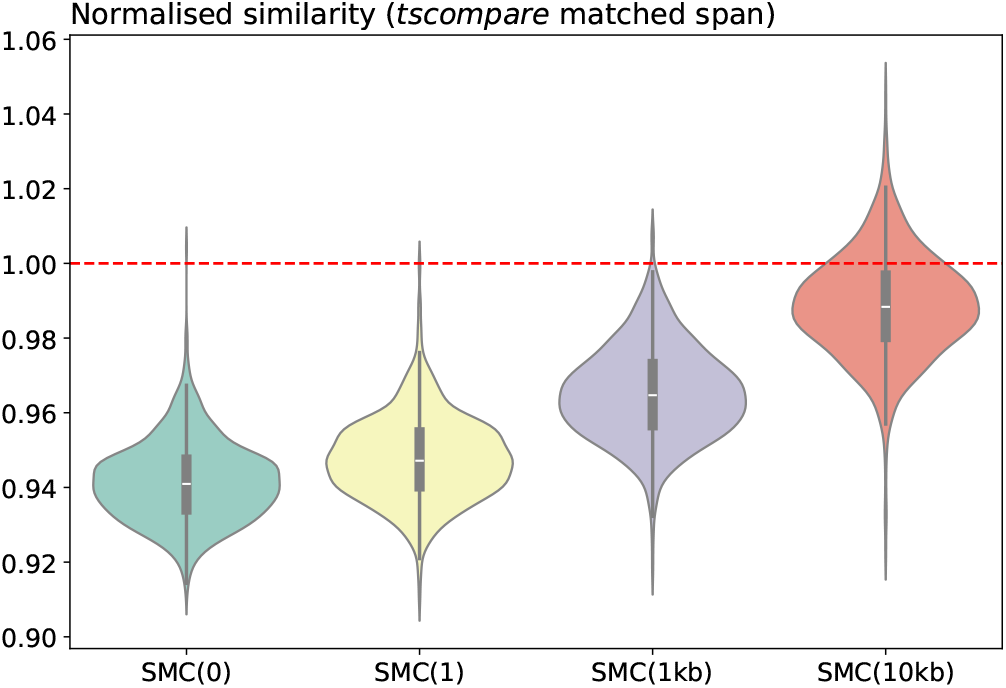
The similarity between ARGs simulated under the CwR and the SMC(k) depends on *k*. The distribution of normalised similarity between ARGs simulated under the CwR and SMC(k) for different values of *k*. Similarity is measured in terms of the `matched span` length computed by *tscompare* normalised by the similarity expected between two random ARGs simulated under the CwR.

### Bootstrapping demographic inference

Drawing inferences about the demographic and selective history of populations is a key goal in population genomics. Many inference tools focus on variation at the level of SNPs which can be summarised via the site frequency spectrum (SFS) or higher order summaries thereof (Gutenkunst *et al*. 2009; Excoffier *et al*. 2021; Liu and Fu 2020; Omarjee *et al*. 2026). While SFS based inference of point estimates in a composite likelihood framework is efficient, it ignores LD between variants. Similarly, inference methods based on variation within sequence blocks (Laetsch *et al*. 2023) or loci also tend to assume statistical independence between them. In both cases, quantifying the uncertainty in composite likelihood estimates has proven challenging, and solutions to this problem generally rely on arbitrary distance cut-offs. That is, one assumes that SNPs or sequence blocks separated by some minimal physical or map distance can be treated as (approximately) statistically independent. The alternative is to perform a data bootstrap with a scale of block-wise resampling that is generally not informed by LD.

This use of distance thresholds – albeit common practice in the field – is clearly *ad hoc* and not justifiable given that under the CwR, LD is non-zero even between pairs of loci on opposite ends of a chromosome (Pluzhnikov and Donnelly 1996; Baird 2015). As a consequence, it is generally unclear whether confidence intervals based on such thresholds are conservative or overconfident. A meaningful estimation of uncertainty of composite likelihood parameters must incorporate the statistical non-independence of genomic variation due to LD, and the SMC(k) provides the means to do this even for organisms with high recombination rates.

To demonstrate how simulations under the SMC(k) can be used to achieve this, we perform parametric bootstraps for the inference of a piecewise constant history of *N*_*e*_ change in the South African *D. melanogaster* population (SP) from WGS data (*n* = 38 individuals (Pool *et al*. 2012)) first described by Chen *et al*. (2024). This history was initially inferred from the unfolded SFS of putatively neutral sites on chromosome 2R (which is free of large inversions) using Stairway Plot 2 (Liu and Fu 2020). To perform a parametric bootstrap for the inferred demographic history, we simulated replicate datasets under the inferred history based on SMC(1). Each simulation replicate consisted of 20 haploid sequences of chromosome 2R assuming a sequence length of 25.3 Mbp, a recombination rate of *r*_bp_ = 1.045 *×* 10^−8^ and a mutation rate of *µ* = 5.49 *×* 10^−9^ (both measured per bp and generation). Replicates were summarised in terms of the mutation based unfolded SFS which we computed using the `tskit` function `allele_frequency_spectrum` (Ralph et al. 2020). We used `blockbuster` (Omarjee et al. 2026), an analytic composite likelihood method analogous to (but more efficient than) Stairway Plot 2, to infer a three epoch model of *N*_*e*_ change from the SFS (code available at https://github.com/hossam26644/smckpaper-figures).

We find that, as expected, the uncertainty in estimated *N*_*e*_ was lowest for the middle epoch (blue in Fig. 6). Since low values of *k* underestimate long range LD, one would expect an associated underestimation of the uncertainty in point estimates of *N*_*e*_. However, while our bootstrap analysis shows the expected increase in uncertainty, measured in terms of mean squared error with *k*, we find that this effect is surprisingly small Fig. S4). In other words, the choice of *k* does not affect inference accuracy significantly. This is in contrast to the logarithmic effect sequence length has on estimates of uncertainty (Fig. 6 b) and Supplementary Fig. S6).

**Figure 6.**
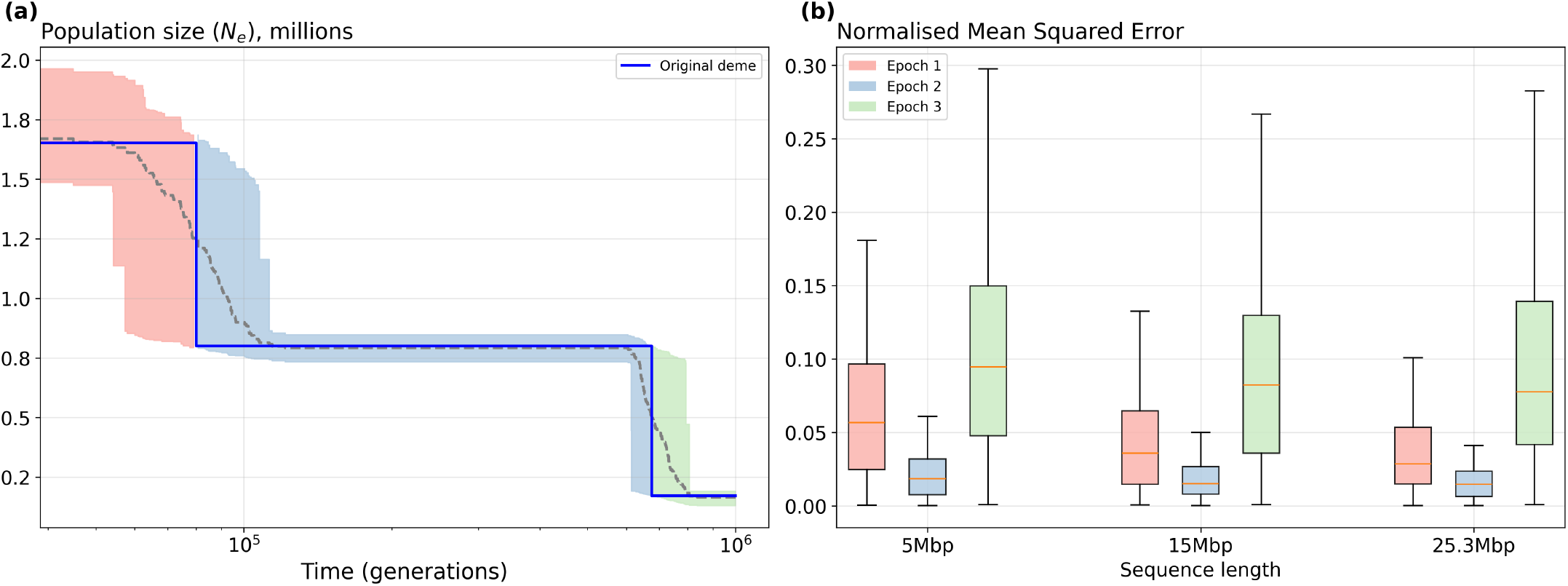
The SMC(k) enables the use of whole chromosome simulations to estimate of the uncertainty in estimates of demographic history. a) The demographic history of the South African *D. melanogaster* population. The blue line represents the piecewise-constant history of *N*_*e*_ change inferred by Chen et al. (2024). The dotted line shows the average history of *N*_*e*_ change inferred using blockbuster across 100 simulated SMC(1) bootstrap replicates; the associated 95% confidence interval is shown as a sleeve. b) The Root Mean Squared Error (RMSE) of epoch wise *N*_*e*_ estimates decreases with sequence length.

This application was enabled by the efficient implementation of SMC(k). For example, a single chromosome-scale bootstrap replicate required approximately 2.6 minutes under SMC(1), giving an estimated total of approximately 4.34 hours for 100 replicates. In contrast, the equivalent CwR simulation would have required approximately 17.6 hours per replicate, or 73.34 days in total.

## Methods

All simulations were performed using *msprime v1*.*4* on an AMD Ryzen™ 9 3950X processor. ARG operations were supported by *tskit v1*.*0* (Jeffery et al. 2026).

### Benchmarking

We averaged the execution time across 10 replicate simulations, each consisting of ten diploid samples of sequence length 25.28 Mbp and a recombination rate *r*_*bp*_ = 1.045*e* − 8. This corresponds to direct estimates of the map length of *Drosophila* chromosome 2R (Comeron et al. 2012, Table 2). We assumed a quadratic fit of the CwR execution time to estimate the predicted execution time for *Drosophila*.

### Comparing SMC(k) and the CwR

We simulated 100kbp of chromosome 2R in *D. melanogaster* (*r*_*bp*_ = 1.045*e* − 8). To measure the distribution of ancestral material, for each of the following parameters, we simulated 10 ARG replicates of two diploid samples with a sequence length of 1Mbp and a population size of 1M diploid individuals, four models: full CwR, SMC(500kb), SMC’ [SMC(1)], and the standard SMC model [SMC(0)]. To retain unary nodes, we set the coalescing_segments_only flag in *msprime* to *false*. Since *msprime* stops simulating a local tree once a most recent common ancestor (MRCA) is reached, unary nodes may be absent from some local trees if they belong to a branch above the local MRCA. Therefore, we set the stop_at_local_mrca flag to false, ensuring that simulations continue until a local MRCA is reached for all local trees. (Fig. 4).

We measured similarity as *tscompare v0*.*2*’s matched sequence span. For different *k* values (0 [SMC], 1 [SMC’], 1kbp, and 10kbp), we simulated 1,000 independent pairs of ARG, each consisting of one ARG simulated under the SMC(k) model and one ARG simulated under the CwR model. Each ARG is made of two diploid samples (Fig. 5).

## Discussion

The central contribution of this work is an efficient implementation of a general and tuneable approximation to the CwR. Unlike previous simulators based on SMC approximations (Chen *et al*. 2009; Staab *et al*. 2015; Shlyakhter *et al*. 2014) our SMC(*k*) implementation is embedded within `msprime`, and can therefore be combined both with its large suite of ancestry models (Baumdicker *et al*. 2022) and flexible demography specifications (Gower *et al*. 2022), and the wider ecosystem of compatible simulation tools (Haller and Messer 2019; Adrion *et al*. 2020; Lauterbur *et al*. 2023; Tsambos *et al*. 2023; Tagami *et al*. 2024). Moreover, `msprime`’s deep integration with `tskit` (Jeffery *et al*. 2026) ensures that the growing suite of efficient ARG operations (Fritze *et al*. 2026; Lehmann *et al*. 2026; Lee *et al*. 2026) can be applied directly to the output. We used these tools to characterise the SMC family of approximations in terms of ARG structure, shedding new light on these well-studied models.

The SMC has usually been evaluated by asking how closely it reproduces summaries of the CwR, in particular standard measures of two-locus LD (Hill and Robertson 1968), the expectation of which can be expressed in terms of the covariance of pairwise coalescence times (McVean 2002). In fact, the consensus that the SMC^*′*^ is a highly accurate approximation of the CwR is entirely based on comparisons of two-locus measures of LD (Paul *et al*. 2011; McVean 2002; Wilton *et al*. 2015). Our results show that this agreement does not extend uniformly to the structure of the ARG as a whole: low-*k* approximations systematically reduce the genomic persistence of ancestral lineages, while increasing *k* progressively restores the long-range ancestral haplotype structure generated by the CwR. We show that this distinction only becomes visible if we move beyond pairwise measures of LD and, instead, consider the full haplotype structure encoded in ARGs directly (Wong *et al*. 2024; Fritze *et al*. 2026).

More generally, our results suggest that there is no single useful definition of the accuracy of an SMC approximation. The SMC^*′*^ approximation reproduces marginal genealogies exactly and approximates two-locus statistics extremely well, while generating ARGS that differ substantially from the CwR in terms of their haplotype structure. The SMC(*k*) provides a convenient way to explore this distinction because *k* defines a controlled sequence of models between the SMC and the CwR.

In practice, the choice of *k* involves a trade-off between computational efficiency and accuracy which must be highly dependent both on the demographic history at hand and the data summary one considers. Our *Drosophila melanogaster* example illustrates this point: increasing *k* has only a modest effect on the estimates of uncertainty for the SFS-based history of *N*_*e*_ we obtain. Thus, reproducing the full ancestral structure of haplotypes under the CwR is clearly unnecessary when quantifying the effect of LD on uncertainty for this particular demographic history. In contrast, the computational gains of SMC^*′*^ make chromosomescale parametric bootstrapping practical in a parameter regime where doing so under the CwR would be prohibitively expensive. It is perhaps unsurprising that *k* matters little when considering histories of *N*_*e*_ change in a single panmictic population.

In contrast, ARGs generated under histories that involve the build up and stochastic decay of long range haplotypes, such as admixture between populations and/or recent selective sweeps may be more sensitive to the choice of *k*. We therefore expect that the SMC(*k*) will be particularly useful for studying the properties of inference methods based on haplotype summaries and reconstructed ARGs for populations that have experienced recent gene flow or selection (Zhang et al. 2026).

The SMC(*k*), and the rich and growing suite of ARG utilities built around `tskit`, allows us to move from asking whether the SMC is “accurate” or not, to asking which properties of ancestry need to be represented in high fidelity for the biological or inferential problem at hand.

## Data availability

## Acknowledgments

We thank Guillaume Achaz for helpful discussions and Stuart JE Baird for comments on an earlier draft.

## Supplementary Information

**Figure S1.**
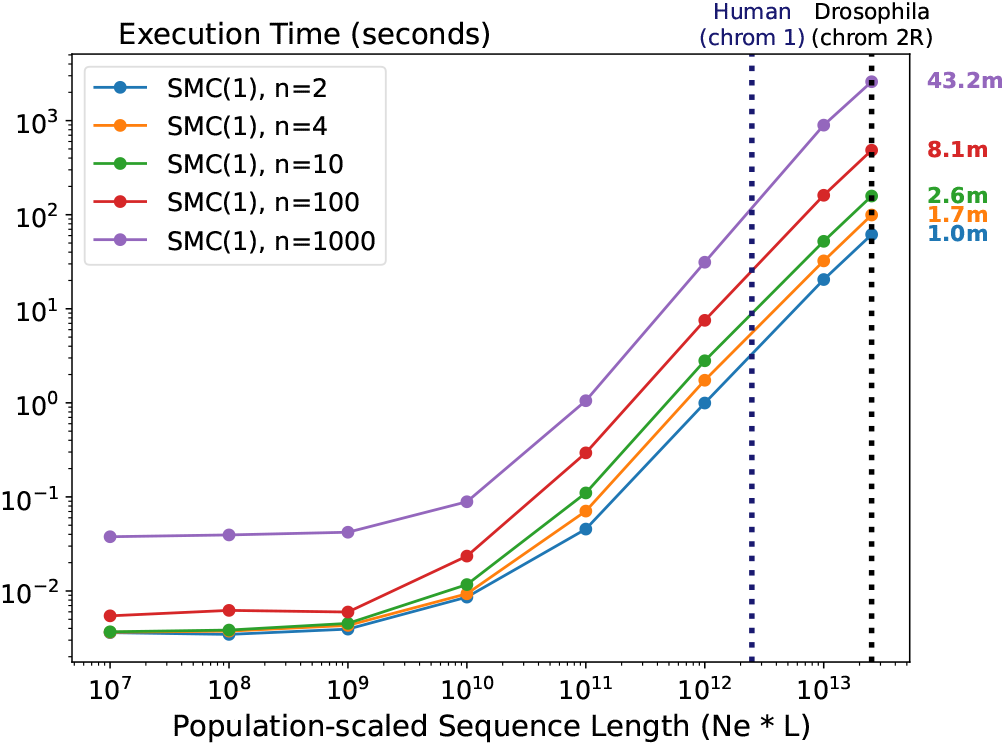
Execution time scales with the number of simulated samples. Execution time in seconds for the SMC(1) model across different population-scaled sequence lengths. Different line colours represent different numbers of simulated samples.

**Figure S2.**
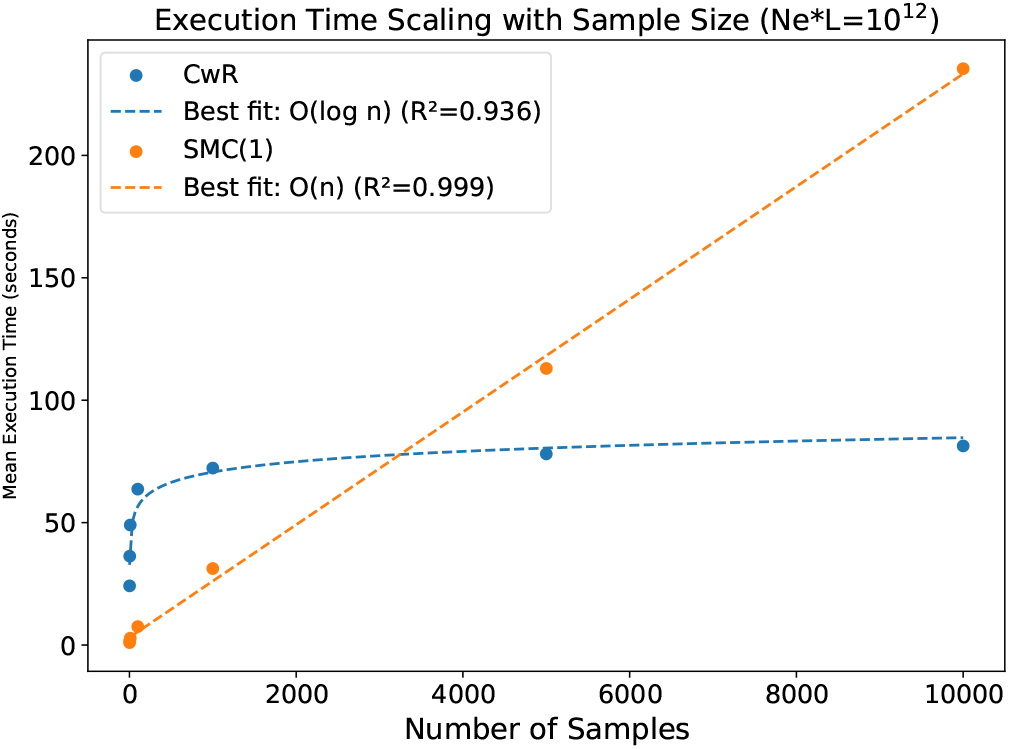
Execution time scales differently between the CwR model and the SMC(1) model. Execution time scales linearly with sample size for the SMC(1) model (*O*(*n*), orange), while it scales logarithmically with sample size for the CwR model (*O*(*logn*), blue).

**Figure S3.**
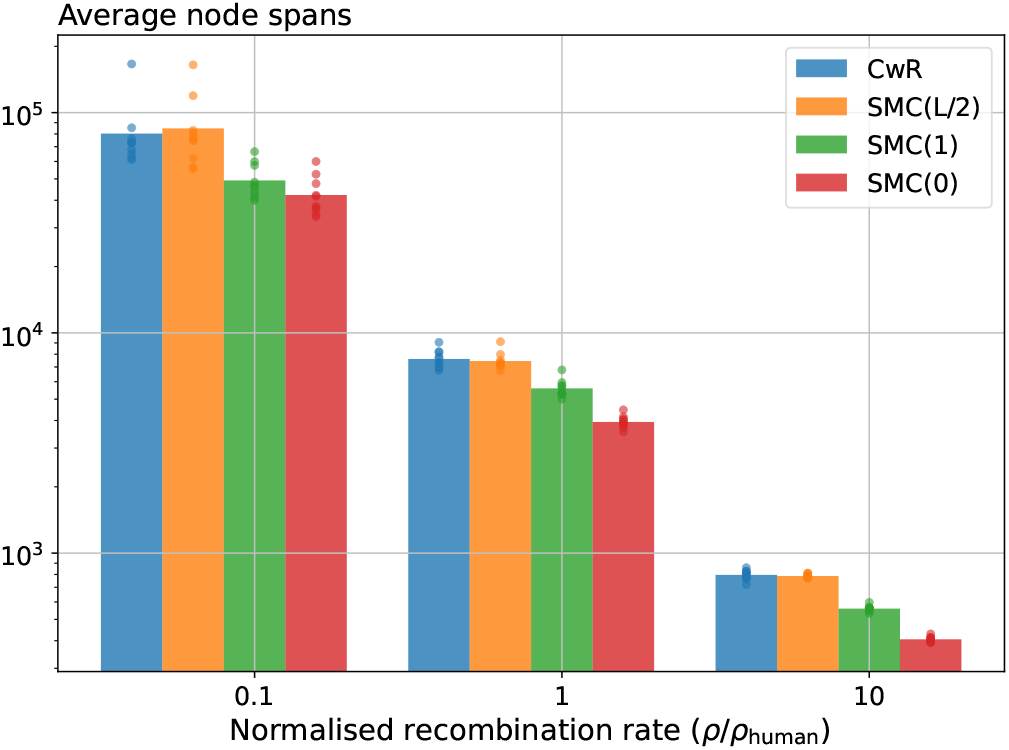
Average binary node spans are shorter for SMC(k) ARGs than for CwR ARGs. The bar plot shows the average binary node span (n=10) for ARGs simulated under different models. The simulated sequence length is 1Mbp, population size is 10^6^ individuals, and recombination rate is 10^−8^.

**Figure S4.**
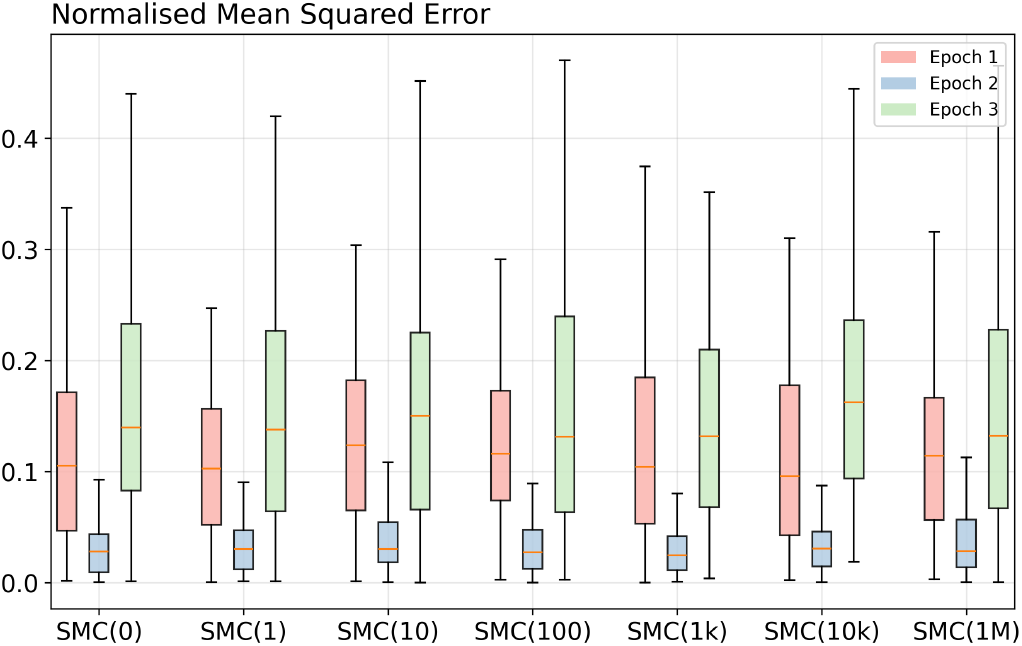
Normalised Mean Squared Error in inferred population size is k value insensitive. *Blockbuster’s* population size inference from SFSs simulated under different SMC models, for a simulated sequence of 1Mbp. Increasing k value does not affect the quality of the population size inference.

**Table S1.** Sequence span per node example. The sequence span per node in Fig:3 categorised by the type of ancestral material.

| Node | Unary Span | Binary Span | Trapped non-ancestral Material Span |
| --- | --- | --- | --- |
| 3 | 5 | 0 | 0 |
| 4 | 4 | 0 | 0 |
| 5 | 3 | 0 | 2 |
| 6 | 3 | 2 | 0 |
| 7 | 2 | 3 | 0 |
| 8 | 0 | 5 | 0 |

**Figure S5.**
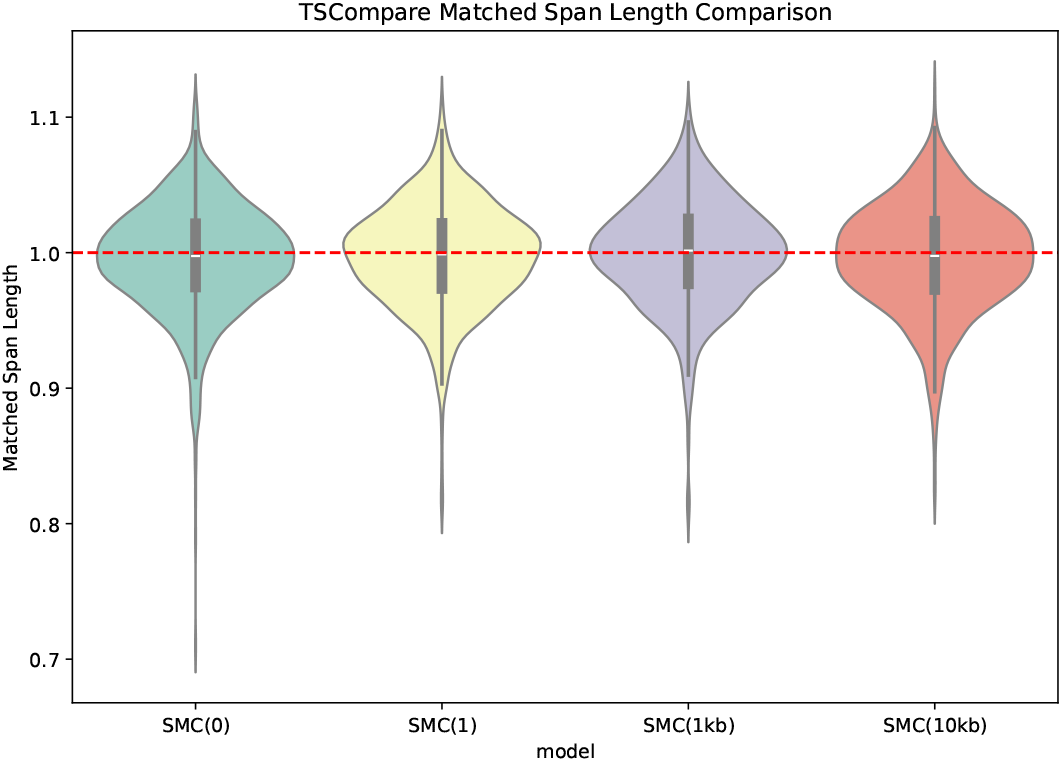
The matched sequence spans of the SMC(K) ARGs to the CwR ARGs does not depend on *k*. The distribution of normalised similarity between ARGs simulated under the SMC(K) and CwR for different values of *k*. Similarity is measured in terms of the matched span length computed by *tscompare v0*.*2* normalised by the similarity expected between two random ARGs simulated under the CwR.

**Figure S6.**
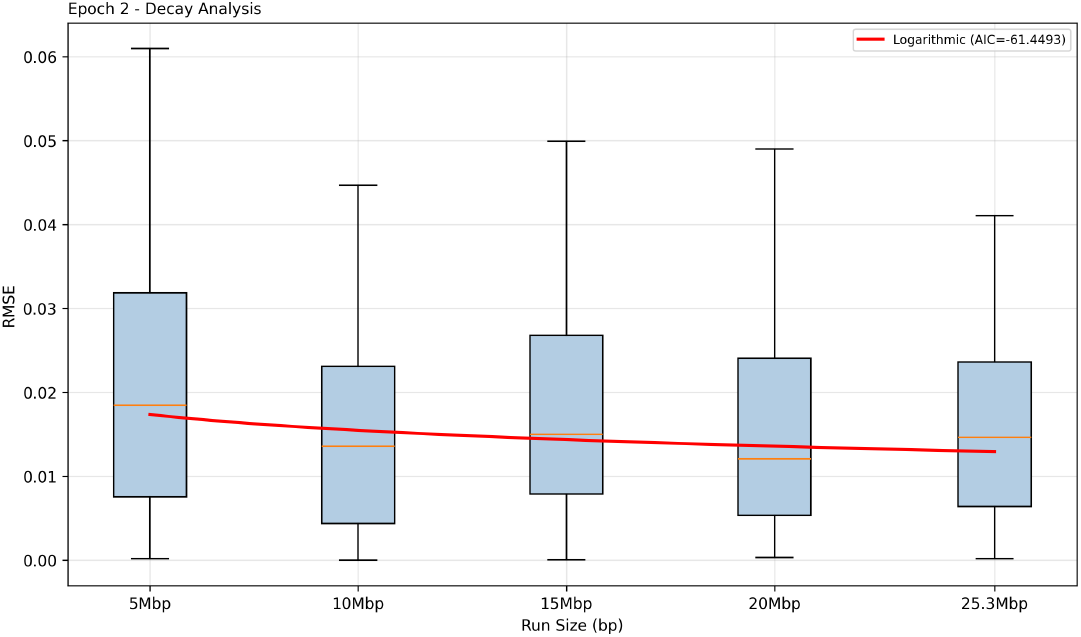
The RMSE error of demographic parameter estimates decreases logarithmically with simulated sequence length.

